# Genetic analysis confirms the presence of *Dicrocoelium dendriticum* in the Himalaya ranges of Pakistan

**DOI:** 10.1101/2020.06.02.130070

**Authors:** Muhammad Asim Khan, Kiran Afshan, Muddassar Nazar, Sabika Firasat, Umer Chaudhry, Neil D. Sargison

## Abstract

Lancet liver flukes of the genus *Dicrocoelium* (Trematoda: Digenea) are recognised parasites of domestic and wild herbivores. The aim of the present study was to address a lack of knowledge of lancet flukes in the Himalaya ranges of Pakistan by characterising *Dicrocoelium* species collected from the Chitral valley. The morphology of 48 flukes belonging to eight host populations was examined in detail and according to published keys, they were identified as either *D. dendriticum* or *Dicrocoelium chinensis*. PCR and sequencing of fragments of ribosomal cistron DNA, and cytochrome oxidase-1 (COX-1) and NADH dehydrogenase-1 (ND-1) mitochondrial DNA from 34, 14 and 3 flukes revealed 10, 4 and 1 unique haplotypes, respectively. Single nucleotide polymorphisms in these haplotypes were used to differentiate between *D. chinensis* and *D. dendriticum*, and confirm the molecular species identity of each of the lancet flukes as *D. dendriticum*. Phylogenetic comparison of the *D. dendriticum* rDNA, COX-1 and ND-1 sequences with those from *D. chinensis, Fasciola hepatica* and *Fasciola gigantica* species was performed to assess within and between species variation and validate the use of species-specific markers for *D. dendriticum*. Genetic variations between *D. dendriticum* populations derived from different locations in the Himalaya ranges of Pakistan illustrate the potential impact of animal movements on gene flow. This work provides a proof of concept for the validation of species-specific *D. dendriticum* markers and is the first molecular confirmation of this parasite species from the Himalaya ranges of Pakistan. The characterisation of this parasite will allow research questions to be addressed on its ecology, biological diversity, and epidemiology.

## 1. Introduction

Digenean lancet liver flukes of the family Dicrocoeliidae (Trematoda: Digenea) can infect the bile ducts of a variety of wild and domesticated mammals and humans around the globe. Three species of the genus *Dicrocoelium*, namely *Dicrocoelium dendriticum, Dicrocoelium hospes* and *Dicrocoelium chinensis* have been described as causes of dicrocoeliosis in domestic and wild ruminants (Otranto and Traversa, 2003). Among these, *D. dendriticum* is the most common and is distributed throughout Europe, Asia, North and South America, Australia, and North Africa (Arias *et al*., 2011). The main economic impact of dicrocoeliosis in livestock is due to the rejection of livers from slaughtered animals at meat inspection (Rojo-Vazquez *et al*., 2012). However, in severe infections, affected animals may show clinical signs including poor food intake, ill thrift, poor milk production, alteration in fecal consistency, photosensitisation and anaemia (Manga-Gonzalez *et al*., 2004; Sargison *et al*., 2012).

The life history of *Dicrocoelium* spp. is indirect and may take at least six months to complete. Monoecious and both sexually reproducing and self-fertilising adults are found in the bile ducts. Eggs containing fully-developed miracidia are shed in faeces and must be eaten by land snails before hatching. Miracidia penetrate the gut wall of the snails and undergo asexual replication and development into cercariae, which then escape from the snails in their slime trails, and are eaten by ants. One cercaria migrates to the head of the ant and associates with the sub-oesophageal ganglion; while up to about 50 cercariae encyst in the gaster as metacercariae (Martín-Vega *et al*., 2018). The larval stage that develops in the ant’s head alters its behaviour, making it cling to herbage and increasing the probability of its being eaten by a herbivorous definitive host. Unlike *Fasciola* spp., larval flukes migrate to the liver via the biliary tree and develop to adults in the bile ducts (Manga-Gonzalez *et al*., 2001). Several species of land snails and ants are known to be intermediate hosts within the same geographical location (Mitchell *et al*., 2017).

Most cases of dicrocoeliosis are subclinical, hence remain undetected (Ducommun and Pfister, 1991). *Dicrocoelium* spp. cause cholangitis; with the thickened bile ducts appearing as white spots on cut surfaces of the liver (Jithendran and Bhat, 1996). The diagnosis of dicrocoeliosis in live animals is usually based on the identification of characteristic eggs in faecal samples, for example by floatation in saturated zinc sulphate solution (Sargison *et al*., 2012). However, coprological methods only detect patent infections, and their sensitivity may be low. Experimental immunodiagnostic methods have been developed to detect anti-*Dicrocoelium* antibodies (Jithendran *et al*., 1996; Gonzalez-Lanza *et al*., 2000; Broglia *et al*., 2009), but these are not available in the field.

Molecular methods amplifying fragments of the ribosomal cistron or mitochondrial loci DNA have been developed (Rehman *et al*., 2020); but as with all molecular diagnostic tools, these depend on accurate speciation of the reference parasites. There are few studies shown the value of ribosomal cistron or mitochondrial loci DNA for the accurate species differentiation of Dicrocoeliidae genera including *D. dendriticum* and *D. chinensis* with little or no information of *D. hospes* (Maurelli *et al*., 2007; Otranto *et al*., 2007; Martinez-Ibeas *et al*., 2011; Bian *et al*., 2015; Gorjipoor *et al*., 2015). These molecular methods have also been adapted to demonstrate the genetic variability and phylogeny of various trematode parasite species affecting ruminant livestock (Chaudhry *et al*., 2016; Ali *et al*., 2018; Sargison *et al*., 2019; Rehman *et al*., 2020) with limited information on Dicrocoeliidae genera.

In this study, we describe the morphological features of *Dicrocoelium* spp. infecting sheep in the Himalaya ranges of Pakistan. The fragments of ribosomal DNA, cytochrome oxidase-1 (COX-1) and NADH dehydrogenase-1 (ND-1) mtDNA were amplified to confirm the species identity of *D. dendriticum*. The genomic sequence data were used to describe the genetic variability and phylogenetic relationships of the Pakistani *D. dendriticum* with the limited number of other *Dicrocoelium* spp. for which matching sequence data are available. Finally, the sequence data were compared with those of *Fasciola hepatica* and *Fasciola gigantica* liver flukes that are the important differential clinical and pathological diagnosis for dicrocoeliosis in ruminant livestock in northern Pakistan.

## 2. Materials and Methods

### 2.1. Fluke collection

A total of 144 sheep from the Chitral valley were examined; comprising of 68 from Booni, 26 from Torkhow, 33 from Mastuj and 17 from the Laspoor valley (Fig. 1). Overall, 639 adult flukes (25 to 50 flukes per animal) were obtained from the livers 16 infected sheep, slaughtered at each of the different localities in the Chitral valley. In all cases, the size and gross morphology of the flukes were typical of *Dicrocoelium* spp. The flukes were washed with phosphate-buffered saline (PBS) to remove adherent debris and fixed in 70% ethanol for subsequent morphometric and molecular characterisation.

**Fig. 1:**
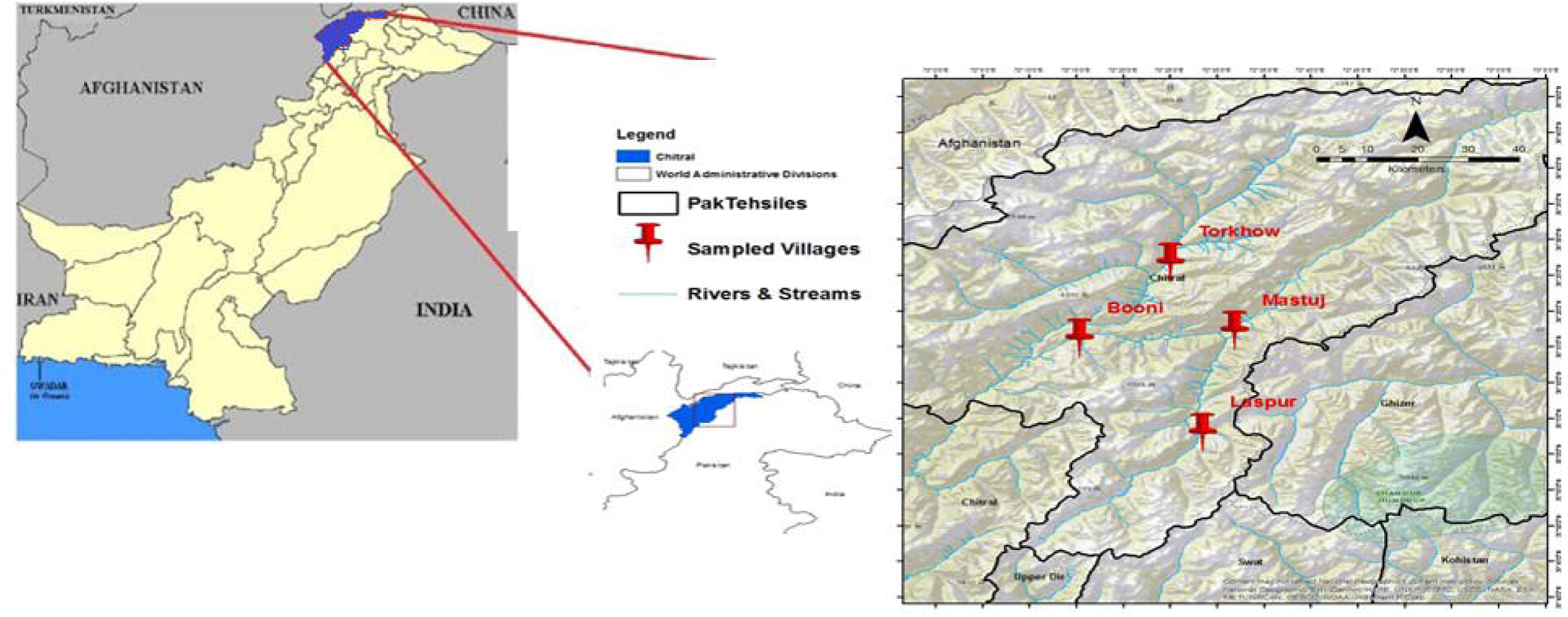
Maps showing the sampling areas in the Chitral valley of the Himalaya ranges of Pakistan.

### 2.2. Morphological examination

Forty-eight adult flukes were selected from the livers of four of 12 sheep from Booni (Pop-1, Pop-2, Pop-3, Pop-4), one sheep from Laspoor (Pop-5), two sheep from Mastuj (Pop-6, Pop-7) and one sheep from Thorkhow (Pop-8), and stained for morphometric analysis. Briefly, the flukes were fixed between two glass slides in formalin-acetic acid alcohol solution, stained with hematoxylin (Sigma-Aldrich) and mounted in Canada balsam. Standardised measurements were taken according to methods described by Taira *et al*. (2006).

### 2.3. PCR amplification and sequence analysis of ribosomal and mitochondrial DNA

Genomic DNA was extracted from 34 individual adult flukes (3 to 6 flukes per animal) from the livers of the same eight sheep (Pop-1 to Pop-8), using a standard phenol-chloroform method (Grimberg *et al*., 1989). A 1,152 bp fragment of the rDNA cistron, comprising of the ITS1, 5.8S, ITS2 and 28S flanking region, was amplified by using two sets of universal primers (Table 1) as previously reported by Gorjipoor *et al*. (2015). A 215 bp fragment of cytochrome c oxidase subunit-1 (COX-1) and a 250 bp fragment of NADH dehydrogenase 1 (ND-1) mtDNA were amplified using newly developed forward and reverse primers (Table 1). Primers were designed with reference sequences of COX-1 and ND-1 of lancet flukes downloaded from NCBI by using the ‘Primers 3’ online tools. The 25 µl PCR reaction mixtures consisted of 2 µl of PCR buffer (1X) (Thermo Fisher Scientific, USA), 2 µl MgCl_2_ (25 mM), 2 µl of 2.5 mM dNTPs, 0.7 µl of primer mix (10 pmol/µl final concentration of each primer), 2 µl of gDNA, and 0.3 µl of Taq DNA polymerase (5 U/µl) (Thermo Fisher Scientific, USA) and 16 µl ddH_2_0. The thermocycling conditions were 96°C for 10 min followed by 35 cycles of 96°C for 1 min, 60.9°C (BD1-F-rDNA/ BD1-R-rDNA), or 61.4°C (Dd-F-rDNA/ Dd-R-rDNA), or 55°C (D-F-cox11/ D-R-cox11), or 57°C (D-F-nad1/ D-R-nad1) for 1 min and 72°C for 1 min, with a final extension of 72°C for 5 min.

**Table 1.**
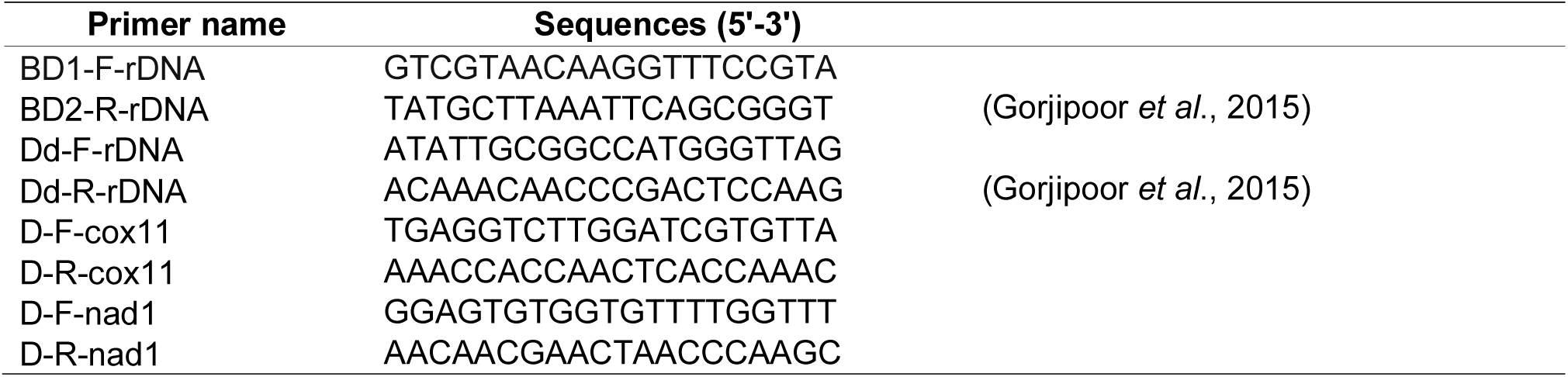
Primer sequences for the amplification of *Dicrocoelium* spp. ITS-2 rDNA, mt-COX-1 mtDNA and ND-1 mtDNA fragments.

PCR products were cleaned using a WizPrepTM Gel/PCR Purification Mini kit (Seongnam 13209, South Korea) and the same amplification primers were used to sequence both strands using an Applied Biosystems 3730Xl genetic analyser. Both strands of rDNA, COX-1 and ND-1 sequences from each fluke were assembled, aligned and edited to remove primers from both ends using Geneious Pro 5.4 software (Drummond, 2012). The sequences were then aligned with previously published NCBI GenBank rDNA, COX-1 and ND-1 sequences of *D. dendriticum, D. chinensis, F. hepatica* and *F. gigantica*. All sequences in the alignment were trimmed based on the length of the shortest sequence available that still contained all the informative sites. Sequences showing 100% base pair similarity were grouped into single haplotypes using the CD-HIT Suite software (Huang *et al*., 2010).

### 2.4. Molecular phylogeny of the rDNA, COX-1 and ND-1 data sets

The generated haplotypes were imported into MEGA 7 (Tamura *et al*., 2013) and used to determine the appropriate model of nucleotide substitution. Molecular phylogeny was reconstructed from the rDNA, COX-1 and ND-1 sequence data by the Maximum Likelihood (ML) method. The evolutionary history was inferred by using the ML method based on the Kimura 2-parameter model for rDNA and Hasegawa-Kishino-Yano model for the COX-1 and ND-1 loci. The bootstrap consensus tree inferred from 1,000 replicates was taken to represent the evolutionary history of the taxa analysed. Branches corresponding to partitions reproduced in less than 50% bootstrap replicates were collapsed. The percentage of replicate trees in which the associated taxa clustered together in the bootstrap test was shown next to the branches. Initial trees for the heuristic search were obtained automatically by applying Neighbor-Join and BioNJ algorithms to a matrix of pairwise distances estimated using the Maximum Composite Likelihood (MCL) approach and then selecting the topology with superior log-likelihood values. All positions containing gaps and missing data were eliminated. There were totals of 698 bp, 187 bp and 217 bp in the final datasets of rDNA, COX-1 and ND-1, respectively. A split tree was created in the SplitTrees4 software by using the UPGMA method in the HKY85 model of substitution. The appropriate model of nucleotide substitutions for UPGMA analysis was selected by using the jModeltest 12.2.0 program.

## 3. Results

### 3.1. Gross liver pathology

Large numbers of flukes were detected in the bile ducts of each liver. The infected livers were indurated and scarred, and the bile ducts were markedly distended, with thickened and fibrosed walls.

### 3.2. Fluke morphometric characteristics

Forty-eight hematoxylin-stained liver flukes were examined. The flukes were all lanceolate, with flattened bodies, and were 5.65 to 8.7 mm in length and 1.3 to 2.2mm in width. The morphology of the examined flukes confirmed the identity of 25 as *D. dendriticum* and 23 as *D. chinensis*, based on the morphology of sub-terminal oral suckers, lobate testes in tandem position behind a powerful acetabulum, small ovaries, ceca simple, a uterus with both ascending and descending limbs and a body is pointed at both ends. The key features of these flukes are shown in Fig. 2 and the morphometric characteristics values are presented in Table 2.

**Table 2:** Range and mean (± standard deviation) of morphometric measurements of *Dicrocoelium* spp. collected from sheep in the Chitral district, Khyber Pakhtunkhwa, Pakistan. The characteristics of 25 flukes were consistent with morphological keys for *D. dendriticum*, and the characteristics of 23 flukes were consistent with morphological keys for *D. chinensis*.

|  | *D. dendriticum* | *D. chinensis* |
| --- | --- | --- |
| **Body** |  |  |
| Length | 5.7 - 8.7 mm ($8.2 \pm 1.0$ mm) | 10.7 - 11.2 mm ($10.9 \pm 0.2$ mm) |
| Width | 1.3 - 2.2 mm ($2.0 \pm 0.2$ mm) | 1.9 - 2.0 mm ($2.0 \pm 0.1$ mm) |
| **Oral Sucker Diameter** |  |  |
| Internal | 150 - 260 μm ($243 \pm 28$ μm) | 210 - 300 μm ($272 \pm 24$ μm) |
| External | 240 - 385 μm ($380 \pm 8$ μm) | 420 - 550 μm ($512 \pm 40$ μm) |
| **Ventral Sucker Diameter** |  |  |
| Internal | 130 - 220 μm ($196 \pm 30$ μm) | 200 - 270 μm ($242 \pm 26$ μm) |
| External | 330 - 410 μm ($390 \pm 24$ μm) | 390 - 420 μm ($402.5 \pm 8$ μm) |
| **Testes** |  |  |
| Length | 575 - 1010 μm ($883 \pm 155$ μm) | 1300 - 1490 μm ($1436 \pm 56$ μm) |
| Width | 500 - 790 μm ($708 \pm 86$ μm) | 695 - 790 μm ($763 \pm 26.2$ μm) |
| **Vitelline glands** |  |  |
| Length | 1.6 - 2.0 mm ($1.82 \pm 0.1$ mm) | 2.1 - 2.3 mm ($2.2 \pm 0.1$ mm) |

**Fig. 2:**
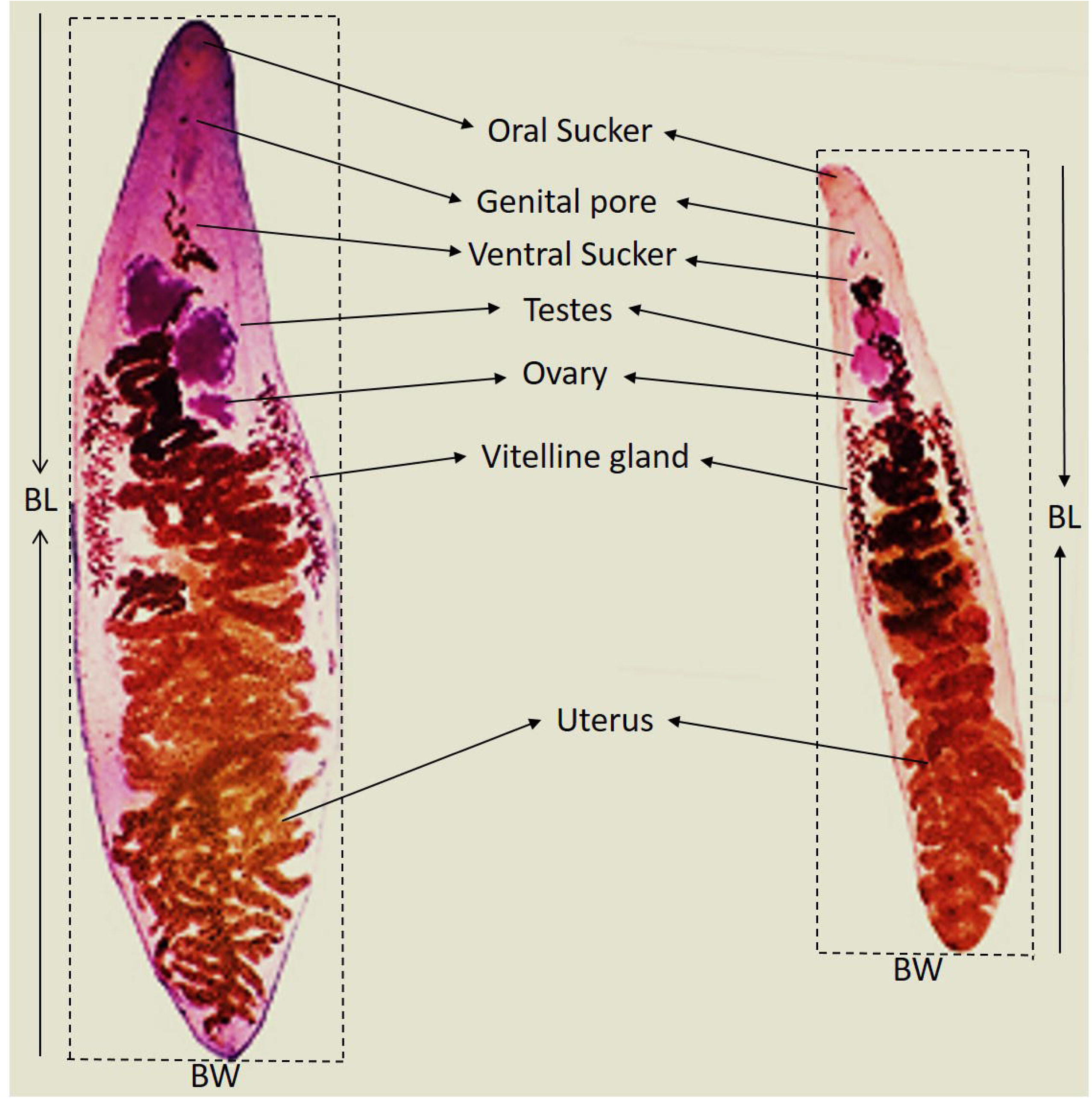
Light micrograph of hematoxylin-stained flukes from the present study. The bodies were pointed at both ends, semi-transparent and pied, with lobate testes in tandem position behind the ventral sucker. The ovary was small, and the black uterus has both ascending and descending limbs and white vitellaria visible under a light microscope.

### 3.3. Molecular confirmation of Dicrocoelium species identity and phylogeny of Dicrocoelium and Fasciola spp

#### 3.3.1. Ribosomal DNA haplotypes

The rDNA sequences of each of 34 flukes were aligned with 12 sequences of *D. dendriticum* and 18 sequences of *D. chinensis* available on the public database (Supplementary Data S1) and trimmed to 698 bp length. This included 4 informative sites to describe intraspecific variation within *D. dendriticum* and 6 sites to describe intraspecific variation within *D. chinensis*. The alignment confirmed 21 species-specific fixed SNPs to differentiate between *D. dendriticum* and *D. chinensis* (Table 3), and allowed the molecular species identity of the 34 flukes to be confirmed as *D. dendriticum*.

**Table 3.** Sequence variation in a 698 bp fragment of rDNA, a 217 bp fragment of ND-1 mtDNA, and a 187 bp fragment of COX-1 mtDNA, differentiating between *D. dendriticum* and *D. chinensis*. The rDNA data are based on 12 sequences identified as *D. dendriticum* and 18 sequences identified as *D. chinensis* on NCBI GeneBank. The ND-1 data are based on 46 sequences identified as *D. dendriticum* and 11 sequences identified as *D. chinensis* on NCBI GeneBank. The COX-1 data are based on 56 sequences identified as *D. dendriticum* and 11 sequences identified as *D. chinensis* on NCBI GeneBank.

| rDNA<br>nucleotide<br>position | *D.*<br>*dendriticum* | *D. chinensis* | ND-1<br>nucleotide<br>position | *D.*<br>*dendriticum* | *D. chinensis* | COX-1<br>nucleotide<br>position | *D.*<br>*dendriticum* | *D. chinensis* |
| --- | --- | --- | --- | --- | --- | --- | --- | --- |
| 18 | T/A | T | 3 | T/A | T | 9 | T | A |
| 56 | C | T | 4 | A/G | A | 12 | T | C |
| 58 | G | A | 6 | T | C | 15 | T | C |
| 60 | T | - | 9 | G/A | G | 24 | G | T |
| 82 | T/C | T | 36 | A/G | A | 30 | G | T |
| 119 | G | A | 37 | A | T | 54 | T/C | T |
| 134 | G | T | 39 | T | A | 57 | T/A | T |
| 221 | C | C/A | 45 | T | C | 63 | T | T/C |
| 222 | T | T/A | 50 | G | G/C | 66 | G/A | G |
| 228 | A | A/G | 51 | T | C | 75 | T | A/T |
| 240 | A | A/G | 57 | T | G | 78 | C/T | T |
| 358 | T | C | 66 | G | T | 84 | A | G |
| 367 | G | A | 78 | G | A | 111 | G | A |
| 370 | C | T | 90 | G | A | 114 | C | T |
| 405 | G | A | 96 | G | T | 129 | T | A |
| 423 | C/T | T | 99 | T | A | 132 | A | G |
| 424 | C | T | 103 | G | G/A | 135 | T | A |
| 469 | T | C | 111 | A | T | 138 | T/C | T |
| 489 | T/C | T | 114 | G/A | G | 141 | T | A |
| 535 | T | C | 118 | G/A | T | 143 | T/G | T |
| 541 | A | G | 126 | T/C | T | 177 | C/T | T |
| 550 | T | C | 139 | G/A | G | 183 | T | C/T |
| 571 | G | A | 142 | C/A/G | A |  |  |  |
| 630 | T | A | 150 | A/G | G |  |  |  |
| 632 | G | T | 156 | A/G | G |  |  |  |
| 654 | T | A | 160 | A | G |  |  |  |
| 655 | C | T | 165 | C/T | T |  |  |  |
| 671 | C | C/A | 167 | C/T | C |  |  |  |
| 680 | G | A | 174 | A | G |  |  |  |
| 681 | T | G | 177 | C/T | G |  |  |  |
| 689 | G | A/G | 178 | G | T |  |  |  |
|  |  |  | 180 | G/A | G |  |  |  |
|  |  |  | 187 | A | T |  |  |  |
|  |  |  | 189 | A | G |  |  |  |
|  |  |  | 199 | A | G |  |  |  |
|  |  |  | 200 | T | T/C |  |  |  |
|  |  |  | 204 | C/T | T |  |  |  |
|  |  |  | 208 | T | T/C |  |  |  |
|  |  |  | 216 | A/G | G |  |  |  |

The 12 *D. dendriticum* sequences from NCBI GenBank and 34 *D. dendriticum* rDNA sequences from the present study were examined along with 18 *D. chinensis*, 59 *F. hepatica* and 34 *F. gigantica* sequences from the public database. Sequences showing 100% base pair similarity were grouped into single haplotypes generating 19, 4, 10 and 5 unique *D. dendriticum, D. chinensis, F. hepatica* and *F. gigantica* haplotypes, respectively (Supplementary Data S1). A ML tree was constructed from the 38 rDNA haplotypes to examine the evolutionary relationship between *D. dendriticum* the other liver flukes. The phylogenetic tree indicates four species-specific clades. *D. dendriticum* and *D. chinensis* form discrete species-specific clades, and *F. hepatica* and *F. gigantica* haplotypes are spread across two separate clades (Fig. 3a).

**Fig. 3:**
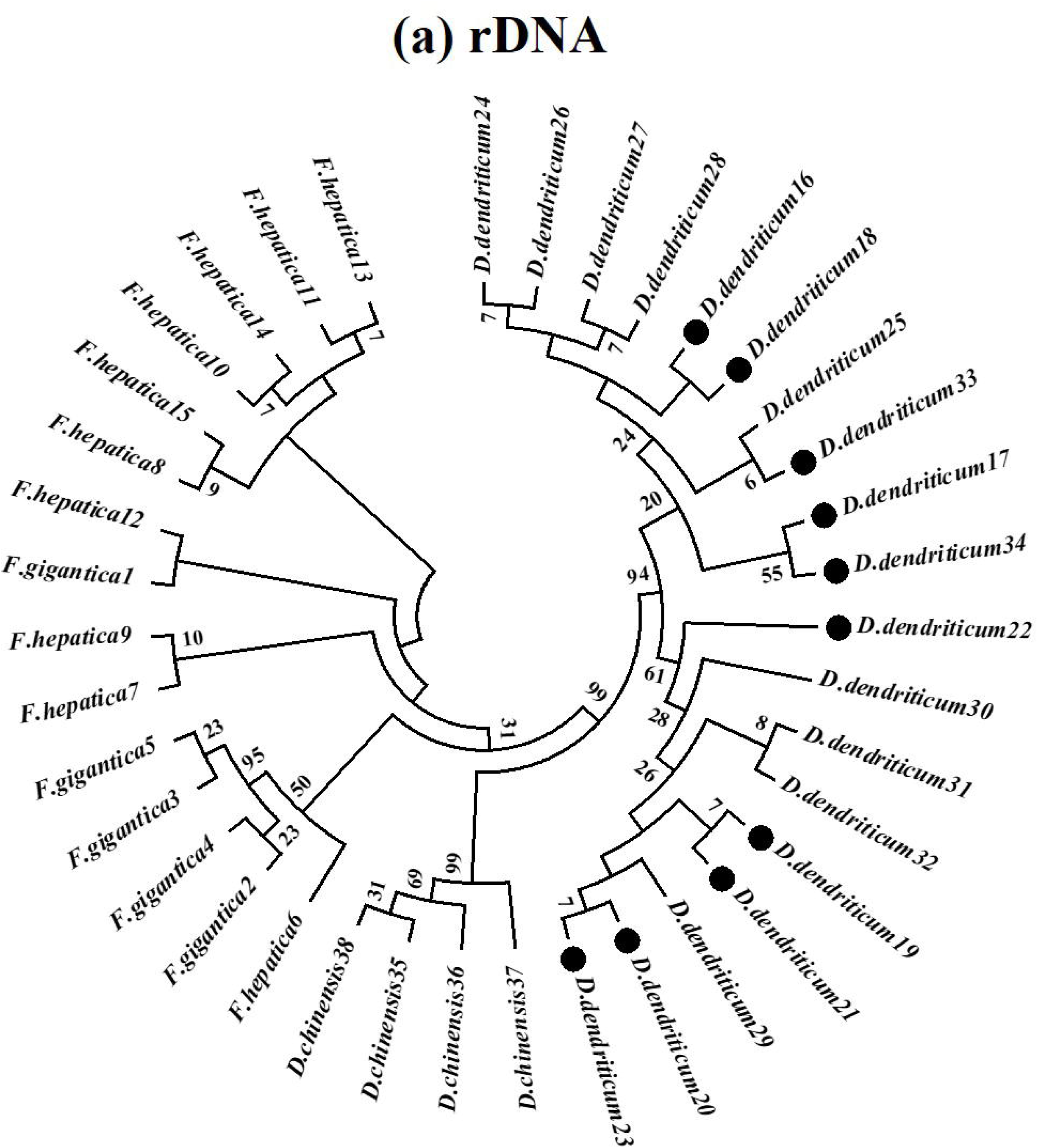
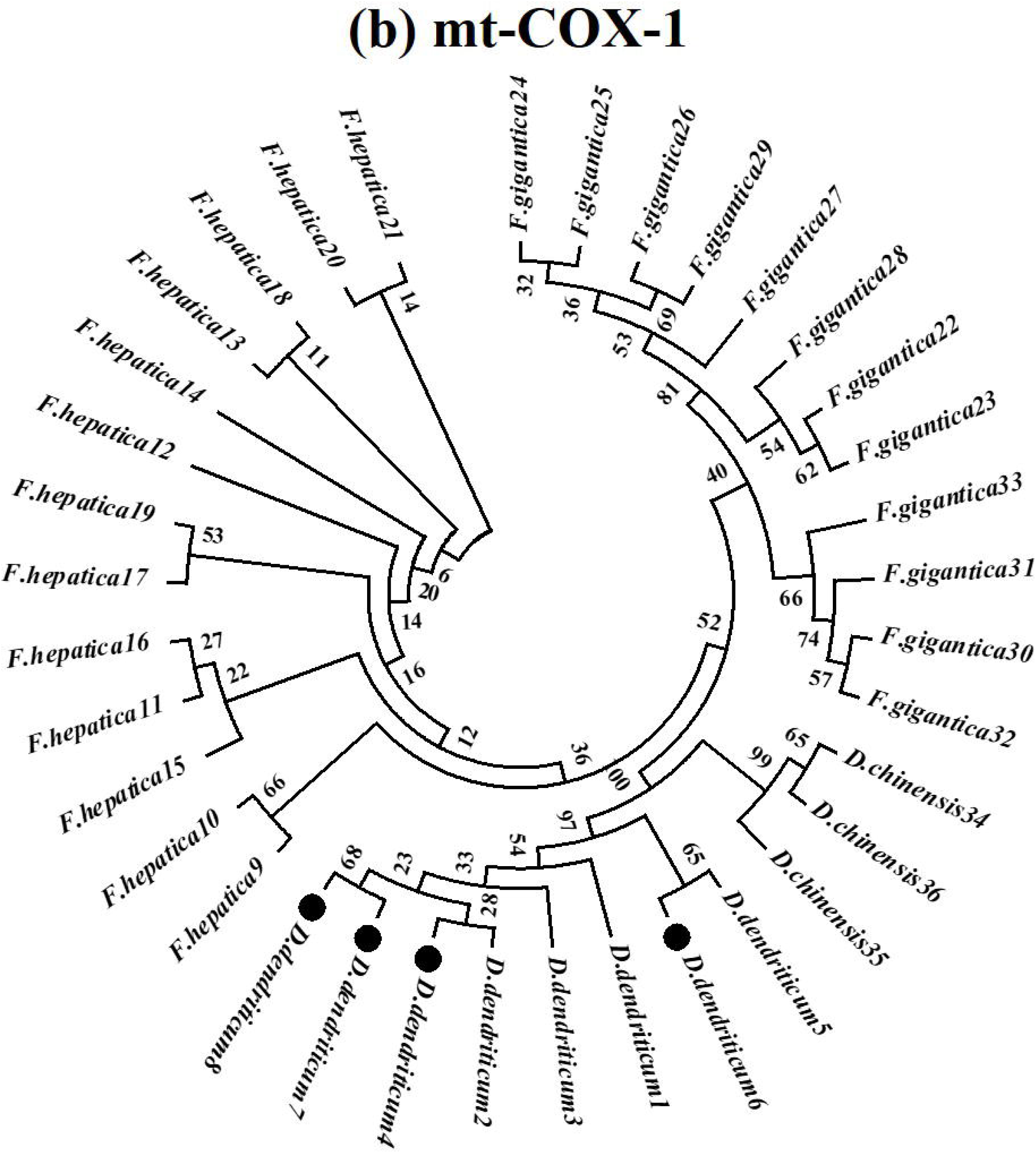
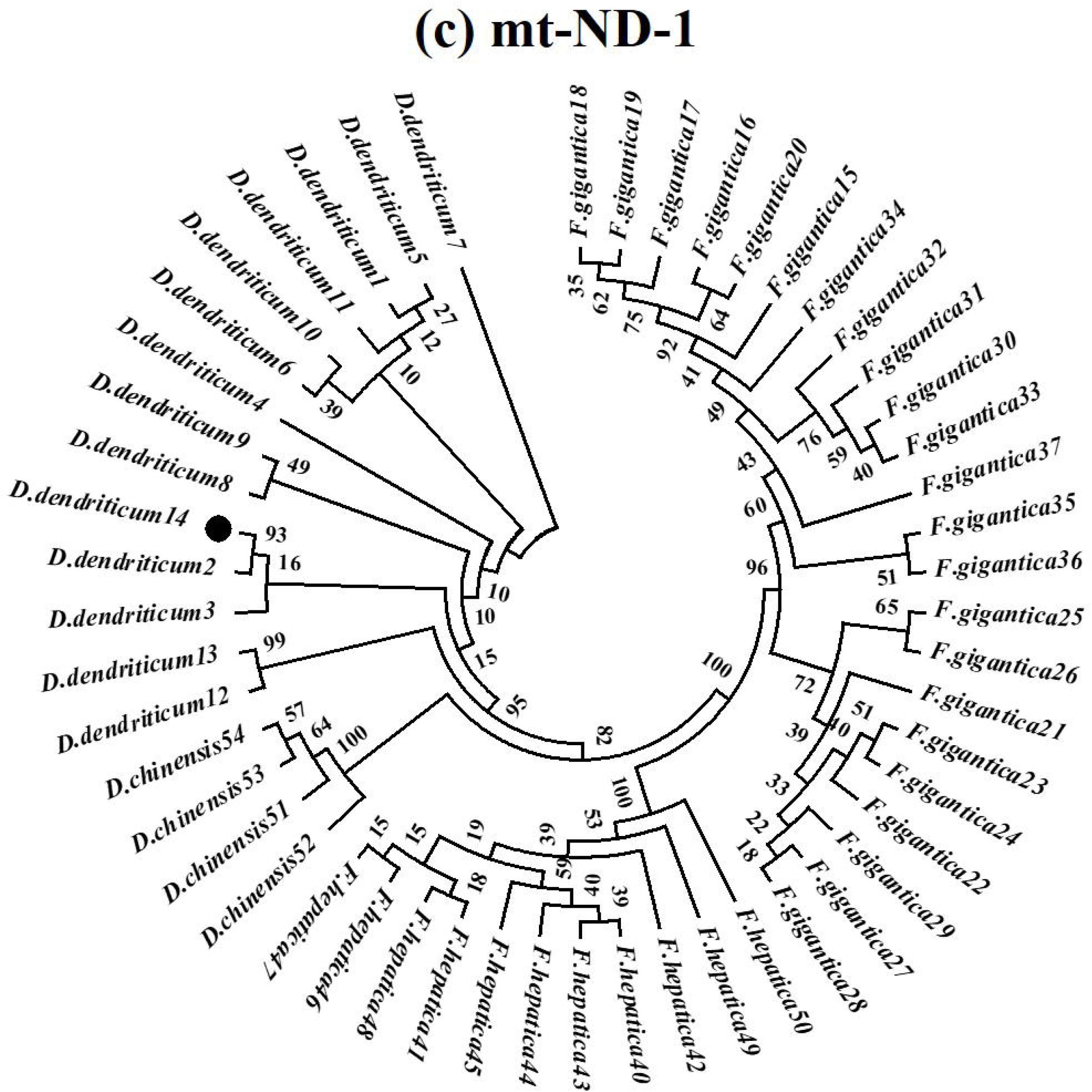
Maximum-likelihood trees were obtained from the rDNA and COX-1 and ND-1 mtDNA sequences of *D. dendriticum, D. chinensis, F. hepatica* and *F. gigantica*. (a) Thirty-eight unique rDNA haplotypes. (b) Thirty-six unique COX-1 mtDNA haplotypes. (c) Fifty-four unique ND-1 mtDNA haplotypes. The haplotype of each species is identified with the name of the species and black circles indicate *D. dendriticum* haplotypes originating from the Chitral valley of Pakistan.

Ten haplotypes generated from the 34 rDNA sequences from the Chitral valley were clustered in the *D. dendriticum* clade (Fig. 3a). Haplotype *D. dendriticum 18* represented two sequences from Pop-1 and Pop-7, which originated from the Booni and Mastuj regions and sequences reported from the Shaanxi province of China. Haplotype *D. dendriticum 16* represented 12 sequences from Pop-2, Pop-3, Pop-4, Pop-6 and Pop-8, which originated from the Booni, Mastuj and Torkhow regions (Fig. 4), while the closely related haplotypes *D. dendriticum 24, 25, 26, 27* and *28* represented sequences from the Shaanxi province of China (Table 4). Haplotype *D. dendriticum 23* represented 4 sequences from Pop-1 and Pop-5 which originated from the Booni and Laspoor regions (Fig. 4), and sequences reported from the Shaanxi province of China. Haplotypes *D. dendriticum 19, 20* and *21* each represented single sequences from Pop-1 and Pop-2 which originated from the Booni region (Fig. 4), while the closely related haplotypes *D. dendriticum 29, 31* and *32* represented sequences from the Shaanxi province of China (Table 4). Haplotype *D. dendriticum 33* represented 9 sequences from Pop-1, Pop-2, Pop-3 and Pop-5 which originated from the Booni and Laspoor regions. Haplotype *D. dendriticum 34* represented 3 sequences from Pop-1 and Pop-6 which originated from the Booni, and Mastuj regions (Fig. 4). Haplotypes *D. dendriticum 17* and *22* represented single sequences from Pop-1 and Pop-6 which originated from the Booni and Mastuj regions (Fig. 4), while the closely related haplotypes *D. dendriticum 25 and 30* represented sequences from the Shaanxi province of China (Table 4).

**Table 4.**
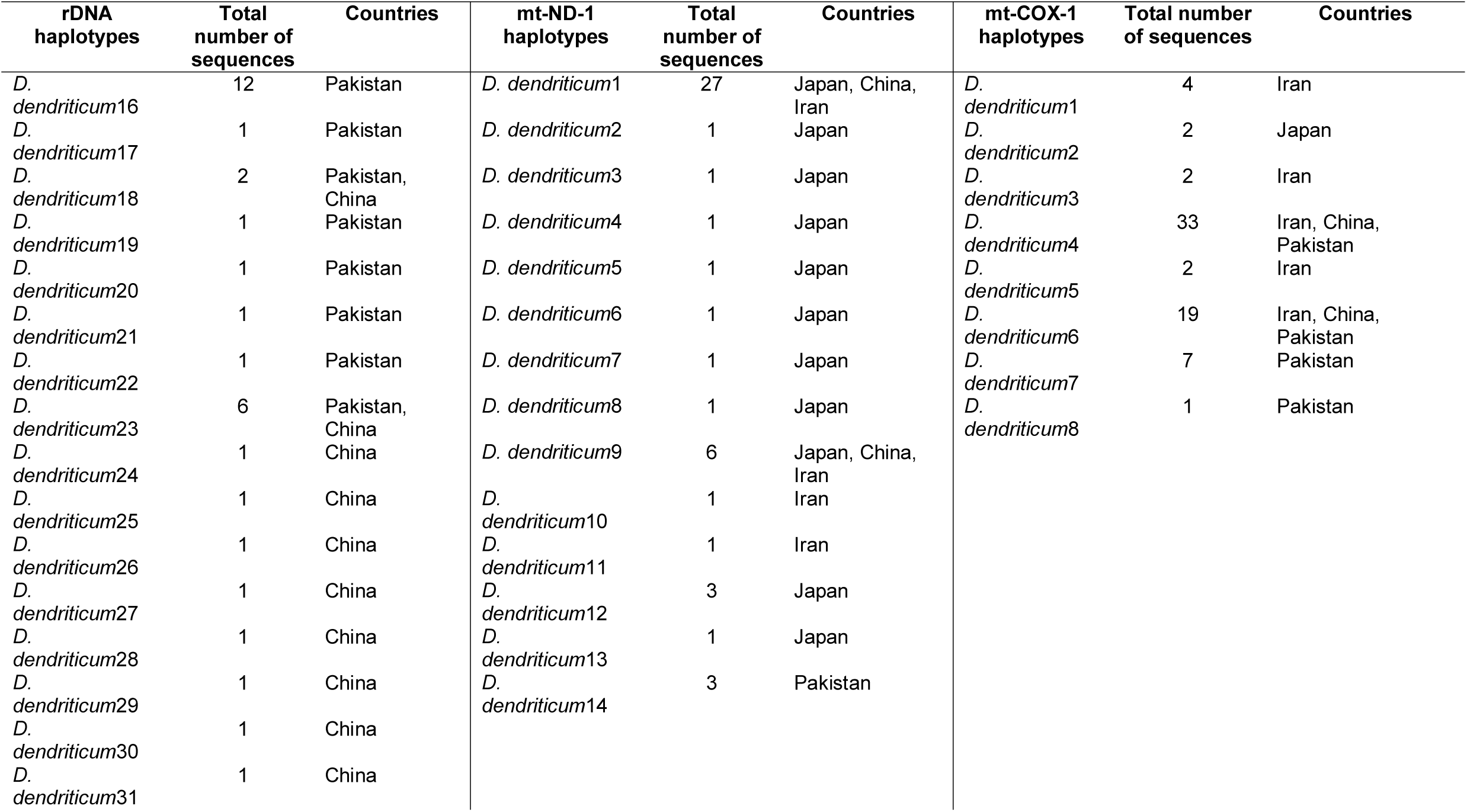
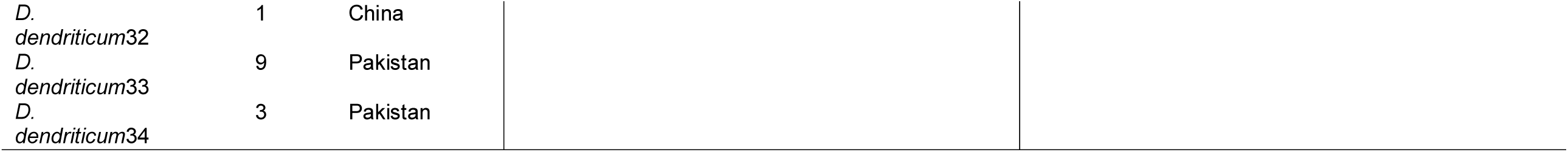
*D. dendriticum* rDNA, ND-1 and COX-1 haplotypes showing the number of sequences representing unique alleles and the country of origin. The accession number of all the sequences are described in Supplementary Data S1, S2 and S3 files.

| rDNA<br>haplotypes | Total<br>number of<br>sequences | Countries | mt-ND-1<br>haplotypes | Total<br>number of<br>sequences | Countries | mt-COX-1<br>haplotypes | Total number<br>of sequences | Countries |
| --- | --- | --- | --- | --- | --- | --- | --- | --- |
| *D. dendriticum*16 | 12 | Pakistan | *D. dendriticum*1 | 27 | Japan, China,<br>Iran | *D. dendriticum*1 | 4 | Iran |
| *D. dendriticum*17 | 1 | Pakistan | *D. dendriticum*2 | 1 | Japan | *D. dendriticum*2 | 2 | Japan |
| *D. dendriticum*18 | 2 | Pakistan,<br>China | *D. dendriticum*3 | 1 | Japan | *D. dendriticum*3 | 2 | Iran |
| *D. dendriticum*19 | 1 | Pakistan | *D. dendriticum*4 | 1 | Japan | *D. dendriticum*4 | 33 | Iran, China,<br>Pakistan |
| *D. dendriticum*20 | 1 | Pakistan | *D. dendriticum*5 | 1 | Japan | *D. dendriticum*5 | 2 | Iran |
| *D. dendriticum*21 | 1 | Pakistan | *D. dendriticum*6 | 1 | Japan | *D. dendriticum*6 | 19 | Iran, China,<br>Pakistan |
| *D. dendriticum*22 | 1 | Pakistan | *D. dendriticum*7 | 1 | Japan | *D. dendriticum*7 | 7 | Pakistan |
| *D. dendriticum*23 | 6 | Pakistan,<br>China | *D. dendriticum*8 | 1 | Japan | *D. dendriticum*8 | 1 | Pakistan |
| *D. dendriticum*24 | 1 | China | *D. dendriticum*9 | 6 | Japan, China,<br>Iran |  |  |  |
| *D. dendriticum*25 | 1 | China | *D. dendriticum*10 | 1 | Iran |  |  |  |
| *D. dendriticum*26 | 1 | China | *D. dendriticum*11 | 1 | Iran |  |  |  |
| *D. dendriticum*27 | 1 | China | *D. dendriticum*12 | 3 | Japan |  |  |  |
| *D. dendriticum*28 | 1 | China | *D. dendriticum*13 | 1 | Japan |  |  |  |
| *D. dendriticum*29 | 1 | China | *D. dendriticum*14 | 3 | Pakistan |  |  |  |
| *D. dendriticum*30 | 1 | China |  |  |  |  |  |  |
| *D. dendriticum*31 | 1 | China |  |  |  |  |  |  |

|  |  |  |
| --- | --- | --- |
| *D.*<br>*dendriticum*32 | 1 | China |
| *D.*<br>*dendriticum*33 | 9 | Pakistan |
| *D.*<br>*dendriticum*34 | 3 | Pakistan |

**Fig. 4:**
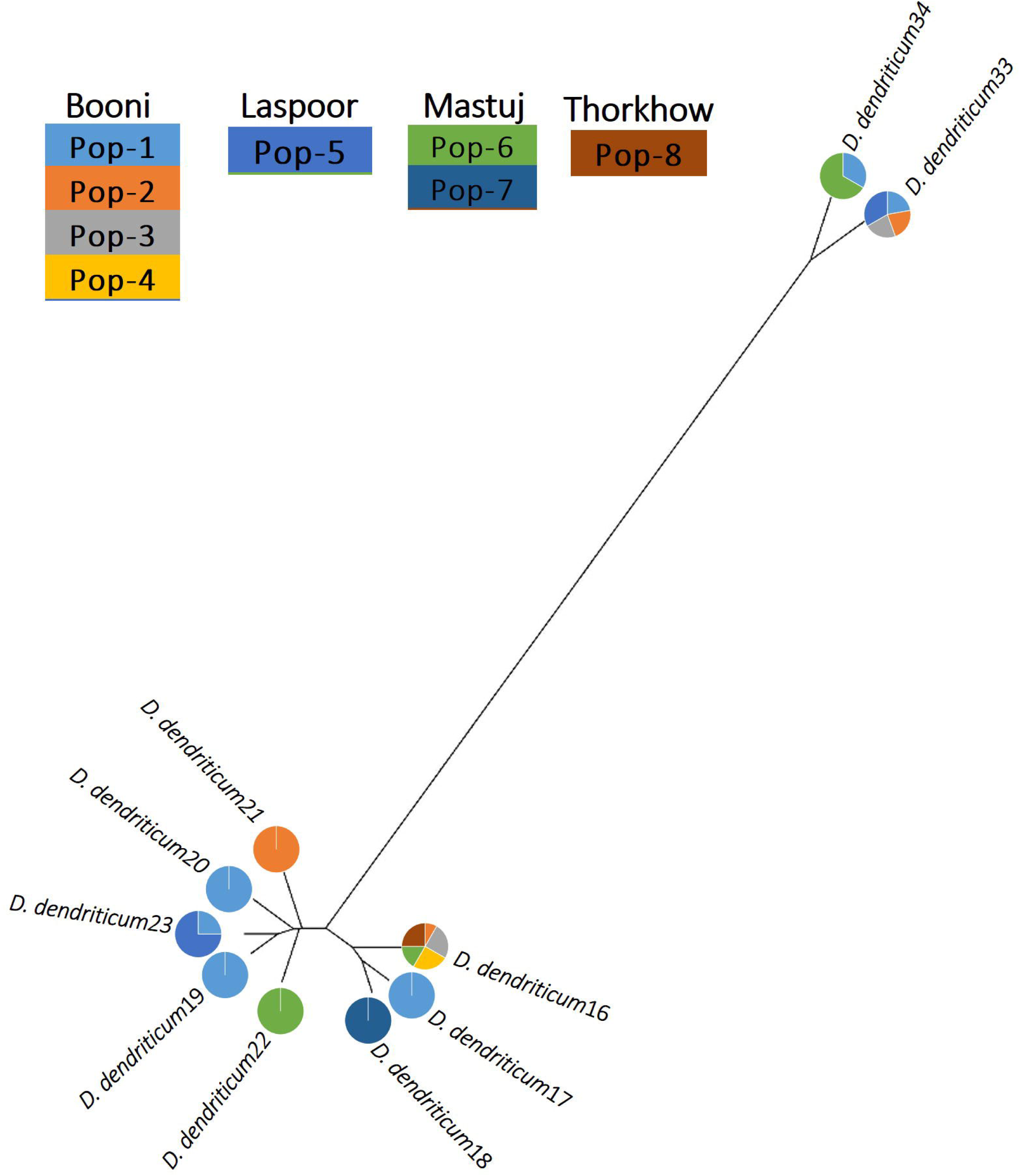
Split tree of 10 rDNA haplotypes generated from eight *D. dendriticum* populations. The tree was constructed with the UPGMA method in the HKY85 model of substitution in the SplitsTrees4 software. The pie chart circle represent the haplotype distribution from each of the eight populations. The color of each haplotype circle represents the percentage of sequence reads generated per population.

#### 3.3.2. Mitochondrial COX-1 haplotypes

COX-1 sequences were generated from 14 flukes from the present study. These were aligned with 56 sequences *D. dendriticum* from NCBI GenBank and 11 sequences of *D. chinensis* available on the public database (Supplementary Data S2) and trimmed to 187 bp length. This included 7 informative sites of intraspecific variation within *D. dendriticum* and 3 sites of intraspecific variation within *D. chinensis*. The alignment confirmed 12 species-specific fixed SNPs to differentiate between *D. dendriticum* and *D. chinensis* (Table 3), allowed the molecular species identity of the 14 flukes to be confirmed as *D. dendriticum*.

The 56 *D. dendriticum* sequences from NCBI GenBank and 14 *D. dendriticum* COX-1 sequences from the present study were examined along with and 11 *D. chinensis*, 99 *F. hepatica* and 83 *F. gigantica* sequences from the public database.

Sequences showing 100% base pair similarity were grouped into single haplotypes generating 8, 3, 13 and 12 unique *D. dendriticum, D. chinensis, F. hepatica* and *F. gigantica* haplotypes, respectively (Supplementary Data S2). A ML tree was constructed from the 36 unique COX-1 haplotypes to examine the evolutionary relationship between *D. dendriticum* and the other liver flukes. The phylogenetic tree indicates four species-specific clades (Fig. 3b).

Four haplotypes generated from the 14 COX-1 sequences from the Chitral valley were clustered in the *D. dendriticum* clade (Fig. 3b). Haplotype *D. dendriticum 4* represented 32 sequences from Pop-2, Pop-3, Pop-5, Pop-6 and Pop-7, which originated from the Booni, Mastuj and Laspoor regions and sequences reported from the Shaanxi province of China and Shiraz province of Iran (Table 4). This haplotype was closely related to *D. dendriticum 2* reported from Japan. Haplotype *D. dendriticum 7* represented sequences from Pop-1, Pop-4, Pop-5, Pop-6 and Pop-8, which originated from the Booni, Mastuj, Laspoor and Torkhow regions. Haplotype *D. dendriticum 8* represented one sequence from Pop-3, which originated from the Booni regions. Haplotype *D. dendriticum 6* represented 17 sequences from Pop-1, Pop-5 and Pop-6 which originated from the Booni, Mastuj and Laspoor regions and sequences reported from the Shaanxi province of China and Shiraz province of Iran (Table 4).

#### 3.3.3. Mitochondrial ND-1 haplotypes

ND-1 sequences were generated from only 3 flukes from the present study. These were aligned with 46 sequences *D. dendriticum* from NCBI GenBank and 11 sequences of *D. chinensis* available on the public database (Supplementary Data S3) and trimmed to 217 bp length. This included 17 informative sites of intraspecific variation within *D. dendriticum* and 4 sites of intraspecific variation within *D. chinensis*. The alignment confirmed 19 species-specific fixed SNPs to differentiate between *D. dendriticum* and *D. chinensis* (Table 3), allowed the molecular species identity of the 3 flukes to be confirmed as *D. dendriticum*.

The 46 *D. dendriticum* sequences from NCBI GenBank and 3 *D. dendriticum* ND-1 sequences from the present study were examined along with and 11 *D. chinensis*, 100 *F. hepatica* and 31 *F. gigantica* sequences from the public database. Sequences showing 100% base pair similarity were grouped into single haplotypes generating 14, 4, 11 and 23 unique *D. dendriticum, D. chinensis, F. hepatica* and *F. gigantica* haplotypes, respectively (Supplementary Data S3). A ML tree was constructed from the 52 ND-1 haplotypes to examine the evolutionary relationship between *D. dendriticum* the other liver flukes. The phylogenetic tree indicates four species-specific clades (Fig. 3c).

One haplotype generated from the 3 ND-1 sequences from the Chitral valley was in the *D. dendriticum* clade (Fig. 3c). Haplotype *D. dendriticum 14* represented sequences from Pop-5, Pop-6, and Pop-8, which originated from the Laspoor, Mastuj and Thorknow regions, while the closely related haplotypes *D. dendriticum 2* and *3* represented sequences from China (Table 4).

## 4. Discussion

The Chitral valley is located in the Himalaya range of Pakistan, north of the Indus River, which originates close to the holy mountain of Kailash in Western Tibet, marking the ranges’ true western frontier. The district has economic and strategic importance as a route of human and animal movements to and from south-east Asia through what is known as the China-Pakistan economic corridor. The average elevation is 1,500 m and the daily mean temperature ranges from 4.1°C to 15.6°C, creating an arid environment with only patchy coniferous tree cover, and providing habitats that are mostly hostile to many snail species. Moving north from Peshawar over the Lowari Pass into Chitral valley the snail fauna changes completely (Naggs *et al*., 2000), potentially creating isolated habitats for *Dicrocoelium* spp. flukes in this region as compared to other parts of Pakistan.

The economics of livestock production is marginal in this region, hence better understanding of any potential production limiting disease, such as dicrocoeliosis is important. The prevalence of fasciolosis is also high in the region, where suitable habitats for the mud snail intermediate hosts of *Fasciola* spp. are created all year round in margins of monsoon rainfall filled ponds, where animals are taken to drink. While various flukicidal anthelmintic drugs are available for use in the control of fasciolosis, few have high efficacy against all stages of *D. dendriticum* (Sargison *et al*., 2012). Control of dicrocoeliosis in livestock, therefore, depends on evasive grazing management and strategic use of anthelmintic drugs. However, while an obvious management control measure for fasciolosis is to provide livestock with clean piped drinking water, current understanding of where and when *D. dendriticum* metacercarial challenge arises is inadequate to inform management for the control of dicrocoeliosis. Accurate diagnostic tests are, therefore, required to improve our understanding of the epidemiology of dicrocoeliosis in specific regions where livestock are kept.

Previous studies have shown the value of morphological and molecular-based methods for the accurate species differentiation of *D. dendriticum* and *D. chinensis* (Otranto *et al*., 2007; Bian *et al*., 2015; Gorjipoor *et al*., 2015). The specimens collected from the Chitral valley were morphologically identified Dicrocoeliidae genus, based on morphological keys. The testes orientation, overall size, and level of maximum body width (Otranto *et al*., 2007) were consistent with the morphological identity of both *D. dendriticum* and *D. chinensis*. However, our molecular analyses showed that *D. dendriticum* and *D. chinensis* can be differentiated and that the Chitral valley lancet flukes were all *D. dendriticum*. The different morphological measurements may reflect different stages of the flukes at the time of collection, normal biological variation, or errors introduced during processing (Taira *et al*., 2006). These factors highlight the complementary value of molecular methods in fluke species identification. The molecular methods in the present study confirmed for the first time the species identity of *D. dendriticum* lancet flukes collected from abattoirs in the Chitral valley of the Himalaya ranges of Pakistan; albeit *D. dendriticum* has been reported in neighbouring Himalayan India, (Bian et al., 2015; Hayashi et al., 2017), China (Shah and Rehman, 2001; Jithendran et al., 1996) and Iran (Khanjari et al., 2014; Meshgi et al., 2019).

The *D. dendriticum* sequences were examined along with *D. chinensis, F. hepatica* and *F. gigantica* sequences from the public database. The analysis of the ribosomal cistron, mitochondrial COX-1 and ND-1 loci showed consistent intraspecific variations in both *D. dendriticum* and *D. chinensis*; while the rDNA, COX-1 and ND-1 loci of *D. dendriticum* and *D. chinensis* differed at 21, 12 and 19 species-specific SNPs, respectively. This consistent intra and interspecific variation within and between *D. dendriticum* and *D. chinensis* allow practical differentiation of the Dicrocoeliidae family. *D. hospes* was not included in our analyses, because comparable sequence data are unavailable. The ML tree that was generated shows separate clades of *D. dendriticum, D. chinensis, F. hepatica* and *F. gigantica*. Hence in the case of co-infections, molecular differentiation between each of the species is possible (Rehman *et al*., 2020).

Ten haplotypes were identified in the ribosomal cistron fragment and four in the mitochondrial COX-1 sequences from the Chitral valley of Pakistan. Mitochondrial ND-1 haplotypes had to be excluded from the analysis of gene flow, because insufficient DNA sequences were generated. Furthermore, comparable sequence data for European and North American *D. dendriticum* populations were unavailable in the public database for analysis. There were both unique and common haplotypes in each *D. dendriticum* population from the Chitral valley, some of which were also present in populations from China, Iran and Japan. There are insufficient data on which to base firm conclusions, but the data imply both gene flow within the region and from other parts of Asia, putatively associated with livestock movements. Further studies based on next-generation methods as described for *Calicophoron daubneyi* rumen flukes and *F. gigantica* liver flukes (Sargison *et al*., 2019; Rehman *et al*., 2020) are needed to define the gene flow and the role of animal movement in the spread of *D. dendriticum*.

The genetic analysis of *D. dendriticum* has implications for the diagnosis and control of dicrocoeliosis in Pakistan. These findings show the potential for the development of population genetics tools to study the changing epidemiology of this parasite, potentially arising as a consequence of changing management and climatic conditions. A better understanding of the molecular evolutionary biology and phylogenetics of *D. dendriticum* will help to inform novel methods for fluke control that are now needed.

## Acknowledgements

The research work presented in this paper is part of PhD dissertation of Muhammad Asim Khan. This study is supported by internal research funds of Quaid-i-Azam University, Islamabad Pakistan. Work at the Roslin Institute uses facilities funded by the Biotechnology and Biological Sciences Research Council, UK (BBSRC).

